# Vaporous essential oils and isolates restore pyrethroid-treated netting efficacy to *Aedes aegypti* (Diptera: Culicidae)

**DOI:** 10.1101/2022.12.13.520257

**Authors:** Leslie C. Rault, Scott T. O’Neal, Ellis J. Johnson, Troy D. Anderson

## Abstract

Decreasing opportunities for mosquitoes to bite is critical in the reduction of mosquito-borne pathogen transmission, such as *Plasmodium spp*. or dengue, yellow fever, and West Nile viruses. Field-evolved resistance to a large selection of synthetic insecticides is crippling efforts to reduce mosquito populations and new strategies are necessary to sustain the efficacy of commercially available tools. *Aedes aegypti* (L.), among other species, have evolved pyrethroid resistance in the field and the pyrethroid-resistant Puerto Rico (PR) strain is a valuable tool for understanding resistance mechanisms. A previous study showed that pyrethroid efficacy can be enhanced by pre-exposing the PR strain to essential oil vapors before topical application of deltamethrin. Insecticide-treated nets (ITNs) and long-lasting insecticidal nets (LLINs) are current products commercialized for mosquito bite protection, but nets using pyrethroids are losing efficacy in the field due to field-evolved pyrethroid resistance. This study tested essential oils previously identified to affect pyrethroid efficacy, as well as their main isolates, to assess if they can restore the efficacy of pyrethroid-treated LLIN against the PR strain. We show that although amyris (*Amyris balsamifera*) expectedly decreases net efficacy, increased mortality can be obtained after exposure to tagetes (*Tagetes bipinnata*) and cajeput (*Melaleuca cajuputi)* oils, but also after exposure to their isolates, such as dihydro tagetone and ocimene, from 1 h after exposure to the net. This study provides a selection of promising synergists used as vaporous emanations to restore pyrethroid efficacy and counteract field-evolved resistance in mosquitoes.

## Introduction

Vector-borne diseases are responsible for over 1 million global deaths annually, with mosquitoes being the main vectors of pathogens (Caraballo and King 2014, Franklinos et al. 2019, World Health Organization 2022). Some of the most significant mosquito vectors are *Anopheles, Aedes*, and *Culex spp*. (World Health Organization 2022). Due to the challenges involved with treating the actual diseases, mosquito bite prevention is the most effective tool for decreasing mosquito-borne disease incidence (Hemingway et al. 2006). However, most vector mosquito species have evolved widespread resistance to nearly all insecticides classes used to reduce populations in the field (Hemingway and Ranson 2000, Ranson et al. 2011, Vontas et al. 2012). These mosquitoes have developed cuticle thickening that decreases penetration (Wood et al. 2010), mutations that reduce interaction of the insecticides with the target site (Hemingway et al. 1989, Thompson et al. 1993, Brengues et al. 2003, Du et al. 2016), and increased detoxification of insecticides by cytochrome P450 monooxygenases, glutathione *S*-transferases, and esterases (Lumjuan et al. 2005, Strode et al. 2008, Balabanidou et al. 2016, Francis et al. 2017, Rault et al. 2019). It is challenging to discover and develop new targets and chemicals for resistant mosquitoes. Thus, it is important to conserve existing products for reducing mosquito populations and community transmission of pathogens.

One of the most widely-used classes of insecticides against mosquitoes is pyrethroids (Ranson et al. 2011, Soderlund 2012). They act on the voltage-gated sodium channel, and although different classes of pyrethroids elicit different symptoms in insects, they generally cause loss of coordination, twitching, and hyper-excitation of the nervous system that leads to death (Gammon et al. 1981). Among mosquito vector species, *Aedes aegypti* is a vector of multiple viruses causing illnesses in humans, such as dengue, chikungunya, and yellow fever, which makes the management of this species a priority for public health (Powers and Logue 2007, Kyle and Harris 2008, Muktar et al. 2016). The control of *Ae. aegypti* relies mainly on pyrethroids due to their relative safety for humans in insecticide-treated nets, particularly alpha-cyano pyrethroids (i.e., alpha-cypermethrin, cyfluthrin, deltamethrin, lambda-cyhalothrin) and non-cyano pyrethroids (i.e., etofenprox, permethrin) (Zaim et al. 2000, Hougard et al. 2003, Manjarres-Suarez and Olivero-Verbel 2013). Two types of pyrethroid-treated nets are commercially available: insecticide-treated nets (ITNs), which required frequent re-treatment and may have exacerbated resistance in the field, and long-lasting insecticide-treated nets (LLINs) with an extended shelf life (Center for Disease Control and Prevention 2019). Although pyrethroid-treated bed nets are primarily an antimalarial tool (Center for Disease Control and Prevention 2019), they have also been utilized in the management of other mosquitoes, including *Aedes aegypti* (Che-Mendoza et al. 2018, Herrera-Bojórquez et al. 2020) *and Culex spp*. (Dery et al. 2012), as well as other insects (GÖKÇE et al. 2018, Ghosh et al. 2021). However, widespread use of pyrethroids has resulted in high selection pressure that has driven field-evolved resistance to pyrethroids in mosquitoes, including *Ae. aegypti* (Ranson et al. 2010, Vontas et al. 2012, Smith et al. 2016). More specifically, the Puerto Rico (PR) strain of *Ae. aegypti* evolved target site mutations in the voltage-gated sodium channel as well as increased cytochrome P450 expression and activity, which protects the species from pyrethroid toxicity (Estep et al. 2017, Rault et al. 2019). We chose the PR strain of *Ae. aegypti* as a model for our study because it exhibits common mutations and enzymatic modifications associated with pyrethroid resistance, and for its accessibility in the U.S. Additionally, it has been the object of studies on pyrethroid and LLIN efficacy, and management strategies for this species would benefit from the addition of new compounds, such as synergists, in the vector-borne disease management toolbox.

We previously demonstrated that several essential oils, as vapors, can restore susceptibility of *Ae. aegypti* PR strain to topically-applied pyrethroids (O’Neal et al. 2019). Here, we show that key isolates of essential oils, as vapors, can increase the efficacy of deltamethrin-treated nets to *Ae. aegypti* PR strain.

## Materials and Methods

### Mosquitoes

The strain of *Aedes aegypti* mosquitoes used in this study was reared and collected as previously described (O’Neal et al. 2019). Briefly, the strain used was a pyrethroid resistant strain, Puerto Rico (PR, obtained through BEI Resources, NIAID, NIH: *Aedes aegypti* PUERTO RICO, MRA-NR-48830). The rearing of mosquitoes for the experiments described in this communication followed methods previously published (O’Neal et al. 2019). For all assays, test subjects consisted of non-blood fed, 3-5 d old adult female mosquitoes, that were aspirated from rearing cages, then anesthetized on ice in the aspiration vial, and transferred to testing cages and net dishes with forceps.

### Gas chromatography and mass spectrometry (GC-MS) of essential oils of interest

The individual components of amyris, cajuput, and tagetes essential oils were analyzed with GC-MS profiling at the University of Nebraska-Lincoln Nebraska Center for Biotechnology, Proteomics and Metabolomics Core Facility (Lincoln, NE). The VOCs in the oil samples were analyzed using an Agilent 7890B gas chromatograph coupled to an Agilent 5977A mass spectrometer (GC-MS). The pure essential oil samples were diluted 500-fold in hexane. The VOCs were chromatographically separated by a 5% phenyl 95% dimethyl arylene siloxane Agilent HP-5MS capillary column (30 m x 0.25 mm I.D.). The GC oven temperature was initially set at 45°C for 3 minutes, then raised to 250°C at 15 °C/min, and finally held for 10 minutes. The split ratio of 2:1 and a constant carrier gas flow (He, 1.2 mL/min) were used. The injector and transfer line between the GC and MS were held at 230°C and 150°C, respectively. The data was analyzed using MassHunter (Agilent version B.07.00). The retention times (RT) of alkane solutions containing C_8_ – C_20_, spiked in the samples, were used to calculate retention indices (RI) for all the VOCs. Putative identification of the VOCs was based on comparison of their RI values and mass spectra with the National Institute for Standard and Technology (NIST) mass spectral library (Johnson 2021). The complete list of oil components can be found in Table 1. The main compounds identified for cajeput and tagetes, not present in the amyris volatile compound profile, were tested in the exposure experiment.

**Table 1:**
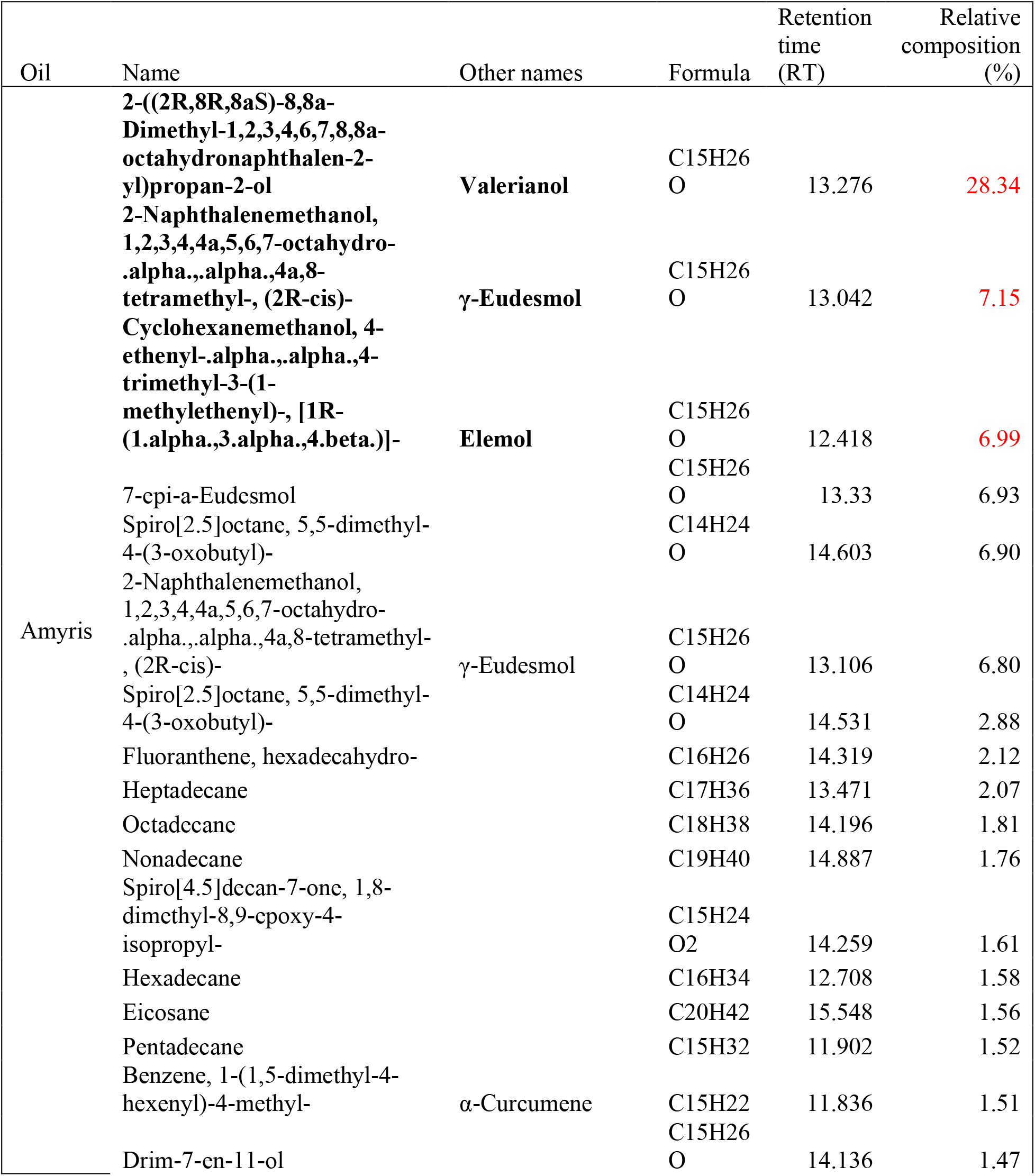
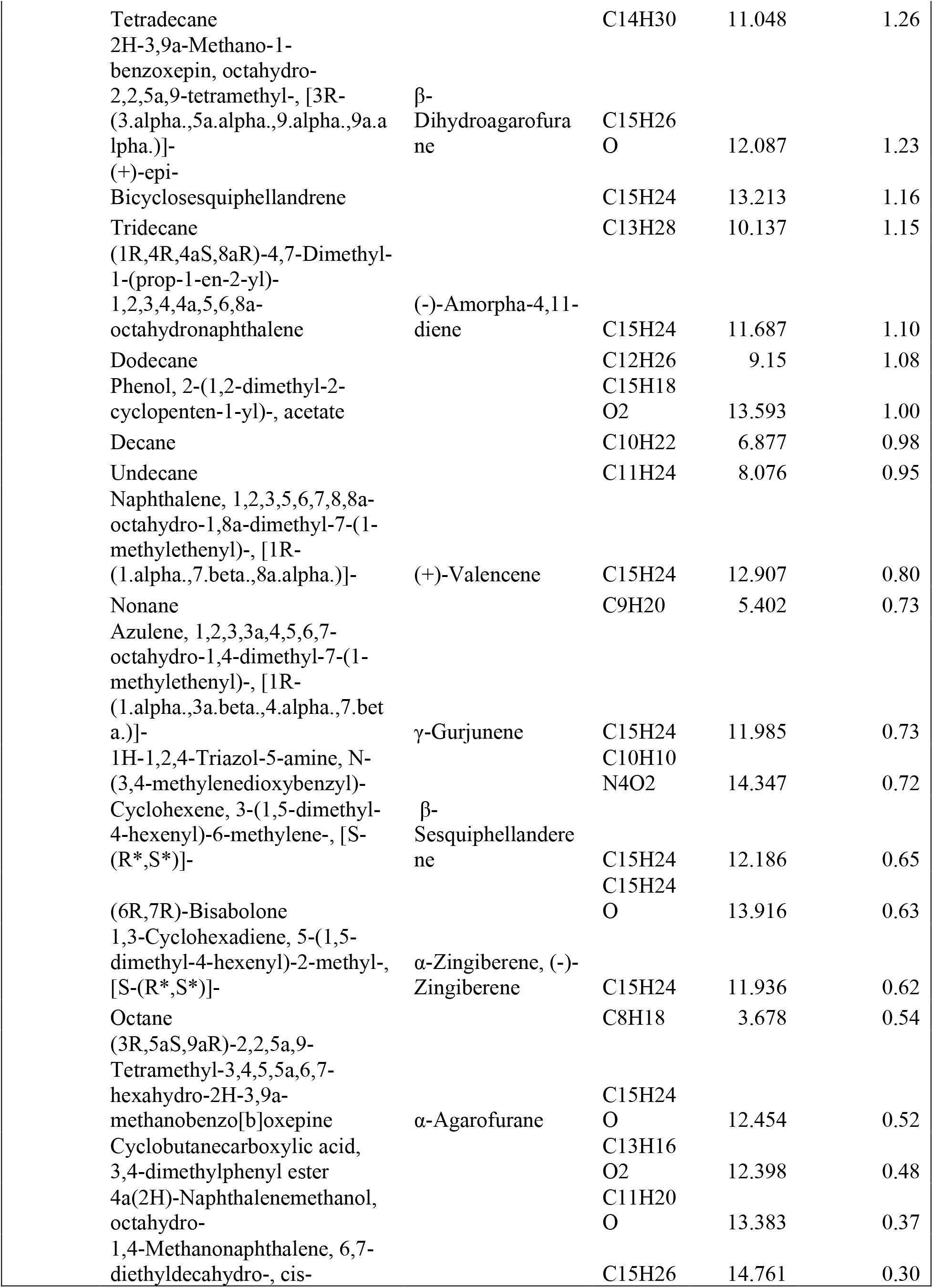
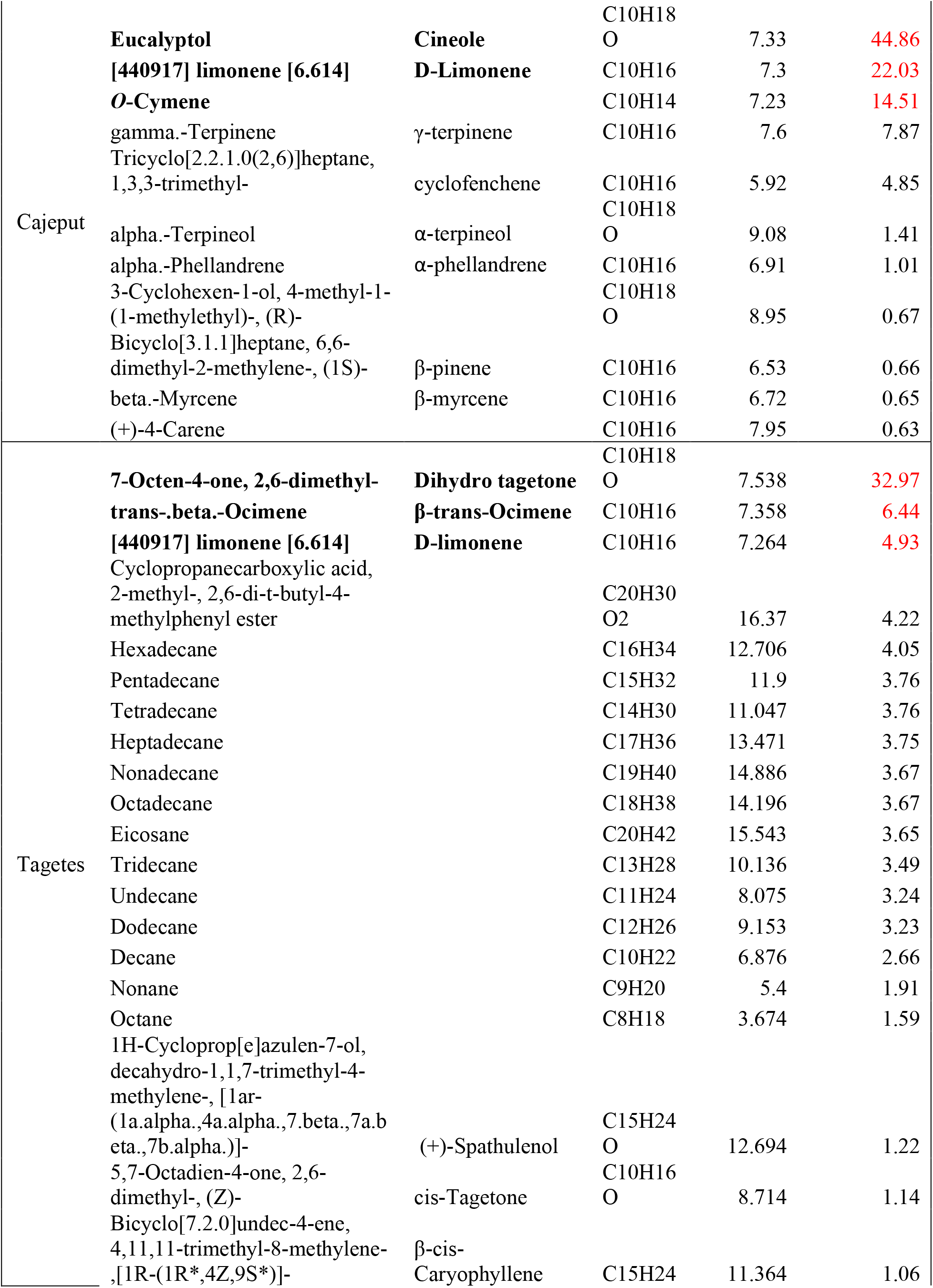
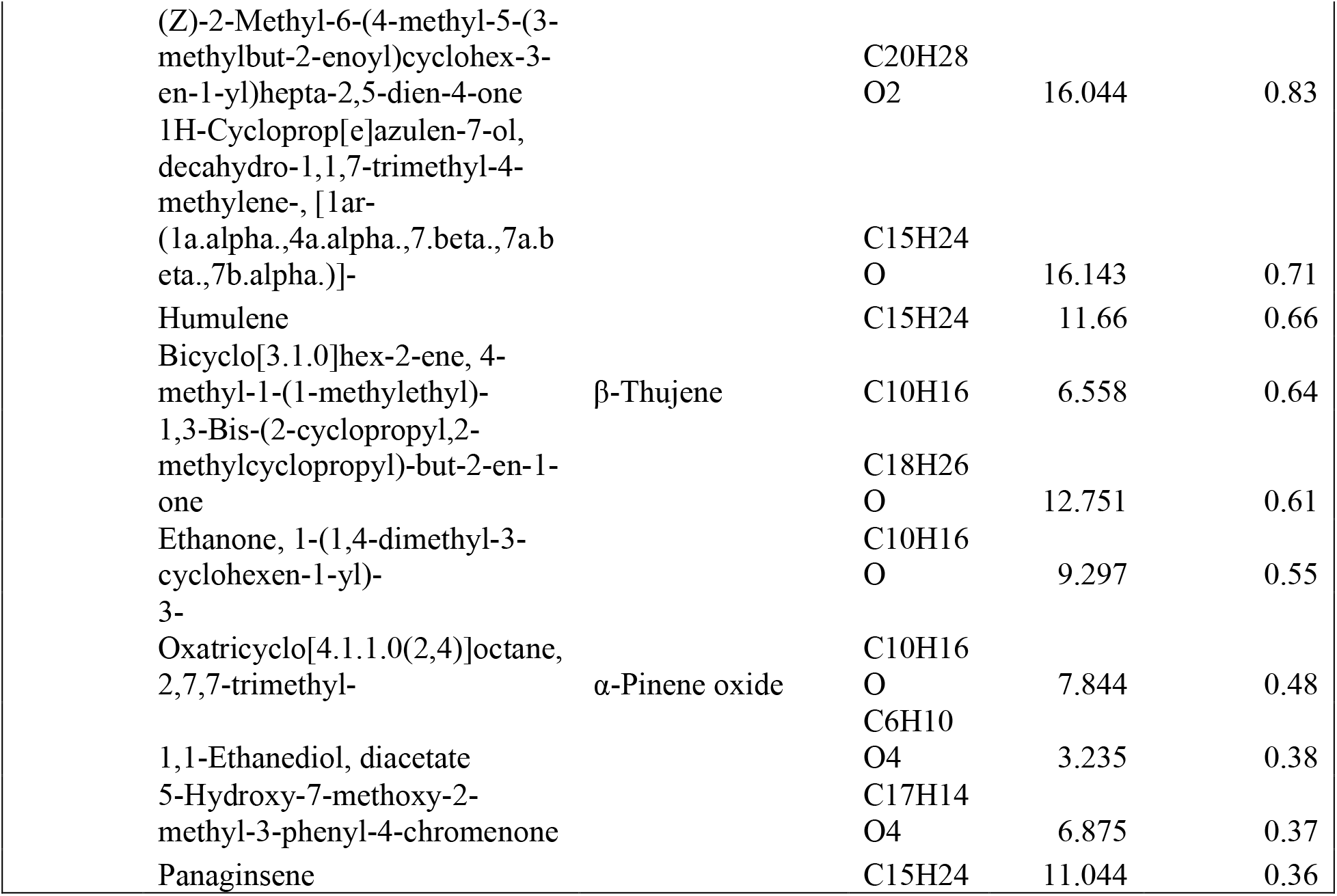
Chemical composition of amyris oil (*Amyris balsamifera*), cajeput oil (*Melaleuca cajuputi*), and tagetes oil (*Tagetes bipinnata*) from GC-MS analysis. In bold are the three most represented compounds in each oil, with their respective percentages in red.

### Treated nets and chemicals

Acetone, >99.5% purity, was purchased from ThermoFisher Scientific (Waltham, MA, USA). PermaNet^®^ screen, a knitted black polyethylene screen/net, or LLIN, containing 0.4% deltamethrin (EPA Reg. No.:85787-1, Vestergaard S.A., Lausanne, Switzerland) was used for the vapor bioassays. A control (untreated) net was used as negative control. Essential oils amyris, cajeput, and tagetes were purchased respectively from BioSource Naturals (Canton, MI), Edens Garden (San Clemente, CA), and Nature’s Kiss (Moreno Valley, CA). Essential oil isolates eucalyptol (C80601, also known as 1,3,3-trimethyl-2-oxabicyclo[2.2.2]octane, 1,8-cineole, 1,8-epoxy-p-menthane) 99%, (*R)*-(+)-limonene, referred to as limonene hereafter (catalog number 183164) 97%, and ocimene (catalog number W353901, mixture of isomers, stabilized) ≥ 90%, as well as hexane for GC-MS dilutions,, were purchased from Sigma (St. Louis, MO) and dihydro tagetone (catalog number sc-391889, 2,6-dimethyl-7-octen-4-one), > 98%, from Santa Cruz Biotechnology (Dallas, TX).

### Vapor bioassays

Vapor bioassays were designed to expose adult mosquitoes to essential oil vapors in an open-air testing environment in order to assess their ability to enhance the contact toxicity of pyrethroid-treated nets, following our previously published method (O’Neal et al. 2019). Ten adult female mosquitoes per replicate were anesthetized on ice and transferred to testing cages (see above). The exposure to undiluted essential oils, isolates, or acetone (as a control) consisted in 100 µL applied to a microcentrifuge cap glued to a Petri dish, and the Petri dish was then placed under the testing cage, that had a perforated bottom to only allow for vapors exposure; no direct contact was possible. Mosquitoes were exposed to the essential oil, isolate, or acetone vapors for 4 h prior to LLIN exposure. Mosquitoes were anesthetized in the testing cup, in the refrigerator (not on ice to avoid having to transfer them to a Petri dish) before being transferred to a Petri dish lined with either control untreated net, or the treated net, for approximately 2 sec. After exposure to the net, mosquitoes were transferred to new testing cups, placed in environmental chambers, and supplied with cotton balls soaked in 10% sucrose until mortality was assessed. Mortality was assessed at the 1 h and 24 h marks following contact with the treated net. Follow-up observations of the 1 h assessments confirmed mortality as opposed to temporary pyrethroid-induced knock-down. Results are reported as mean percent mortality ± standard deviation for each treatment. Mosquito mortality in the controls was also accounted for, and if control mortality exceeded 20%, data from that test group were not used (Abbott 1925). For each treatment or control, 6 to 12 replications were used, all consisting of 10 female mosquitoes.

### Statistical Analysis

All calculations and statistical analyses were performed with GraphPad Prism 9 (GraphPad Software, Inc., La Jolla, CA). Mortality differences for both timepoints together, as repeated measures, were statistically compared to their respective untreated controls (acetone, untreated net) using a three-way ANOVA followed by Tukey’s multiple comparison test (Zar 2010). All results with a *P*-value and an adjusted *P*-value (*P* adj.) of 0.05 or below (≤ 0.05) were considered significant.

## Results

The major components of cajeput (*Melaleuca cajuputi*) oil were eucalyptol, D-limonene, and *o*-cymene at 44.86%, 22.03%, and 14.51% respectively, making up over 80% of cajeput oil. D-limonene was also found in tagetes (*Tagetes bipinnata*) oil, at 4.93% (third most abundant). Dihydro tagetone was the most relatively abundant compound representing 32.98% of the oil, whereas the second most abundant compound was trans-β-ocimene (ocimene) at 6.44% (Table 1). None of these compounds were identified in amyris (*Amyris balsamifera*) oil (Table 1). Only *o*-cymene was not tested in the following experiment.

Based on the Tukey’s multiple comparison (Supplementary data 1) of essential oils and their controls, amyris, cajeput, and tagetes oils alone, delivered as vapor pre-treatments, did not significantly kill more mosquitoes than the acetone control without exposure to deltamethrin after 1 or 24 h (Figure 1) *P* adj. > 0.9999 for all three variations at 1h of 0%, +2.13E-14%, 0% for amyris, cajeput, and tagetes, respectively; at 24 h -1.262%, -6.363%, and +3.03% respectively). Mortality 1 h after exposure to deltamethrin-treated nets was significantly decreased by pre-exposure to amyris compared to the control, *P* adj. < 0.0001, - 30.83%, but significantly increased by pre-exposure to cajeput, *P* adj. = 0.0002, +22.73%, and tagetes, *P* adj. < 0.0001, +29.85% (Figure 1.). After 24 h, these trends are conserved and more specifically, although acetone pretreatment does increase mortality 24 h after exposure to the LLIN (+21.67%, *P* adj. < 0.0001), amyris still decreases mortality by 36.44%, *P* adj. < 0.0001. Cajeput increased mortality by 16.82%, *P* adj. = 0.0047, and tagetes by 35.60%, *P* adj. < 0.0001, reaching 100% mortality (Figure 1).

**Figure 1:**
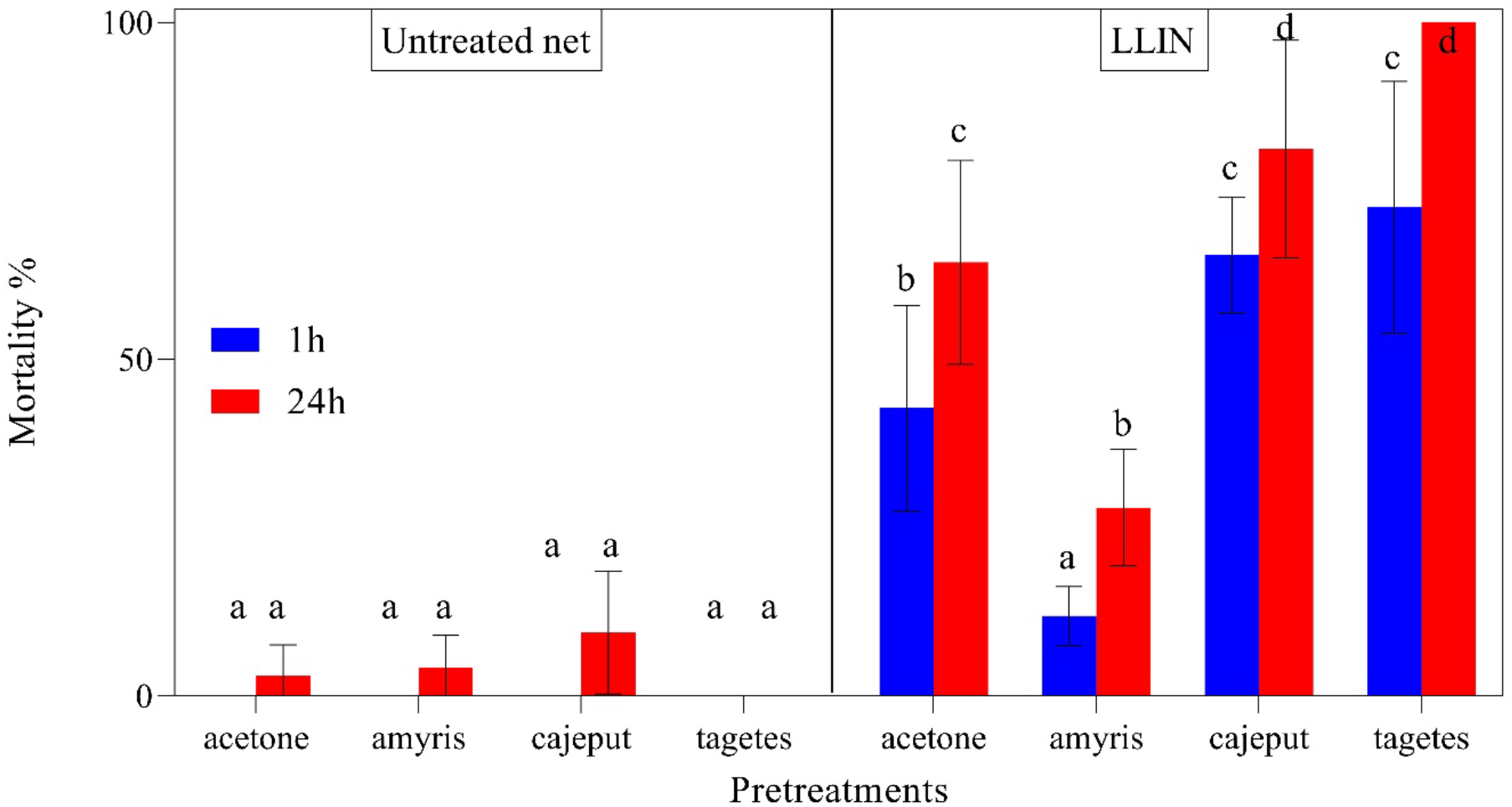
Mortality percentages of 3-5 d old adult female Puerto Rico (*Aedes aegypti*) mosquitoes exposed to essential oil or acetone (control) vapors for 4 h-pre-treatments and transferred briefly to either an untreated net (control, left), or a LLIN (right). Mortality was assessed after 1 h (blue) and 24 h (red) post-contact with the nets. Data presented as mean mortality percentages ± standard error for 60-120 mosquitoes (10 mosquitoes per replicate, number of replicates varies per treatment due to repetition of certain conditions). Statistical differences were obtained with a three-way ANOVA and Tukey’s multiple comparison test, statistical differences are indicated by different letters.

The main isolates identified in these essential oils were dihydro tagetone, eucalyptol, limonene, and ocimene, which were tested as vapors using the same approach. The Tukey’s multiple comparison test (Supplementary data 2) showed that the mortality obtained with vapor exposure to the isolates alone did not differ from the acetone control when untreated by the insecticide net after 1 or 24 h (Figure 2), 0% variability, except eucalyptol with 2.84E-14% increase after 1 h and *P* adj. > 0.9999, and after 24 h +0.3033%, +6.363%, -3.030%, -7.105e-015% and *P* adj. > 0.7537 respectively for dihydro tagetone, eucalyptol, limonene, and ocimene. Mortality 1 h after exposure to deltamethrin-treated nets was significantly increased by pre-exposure to dihydro tagetone by 37.58%, *P* adj. < 0.0001, and ocimene by 27.58%, *P* adj. < 0.0001, but not eucalyptol with an increase of 4.697%, *P* adj. = 0.8248, or limonene at 1.063%, *P* adj. = 0.9990 (Figure 2), compared to the control (acetone-pretreated LLIN-exposed mosquitoes). After 24 h, the trends were conserved as mortality was significantly increased by pre-exposure to dihydro tagetone by 35.60%, *P* adj. < 0.0001, and ocimene by 35.60%, *P* adj. < 0.0001, compared to the 24 h LLIN-exposed acetone pretreated mosquitoes, both dihydro tagetone and ocimene LLIN treatments reaching 100%, but not eucalyptol at 10.91%, *P* adj. = 0.0864 or limonene at 9.698%, *P* adj. = 0.1506 (Figure 2).

**Figure 2:**
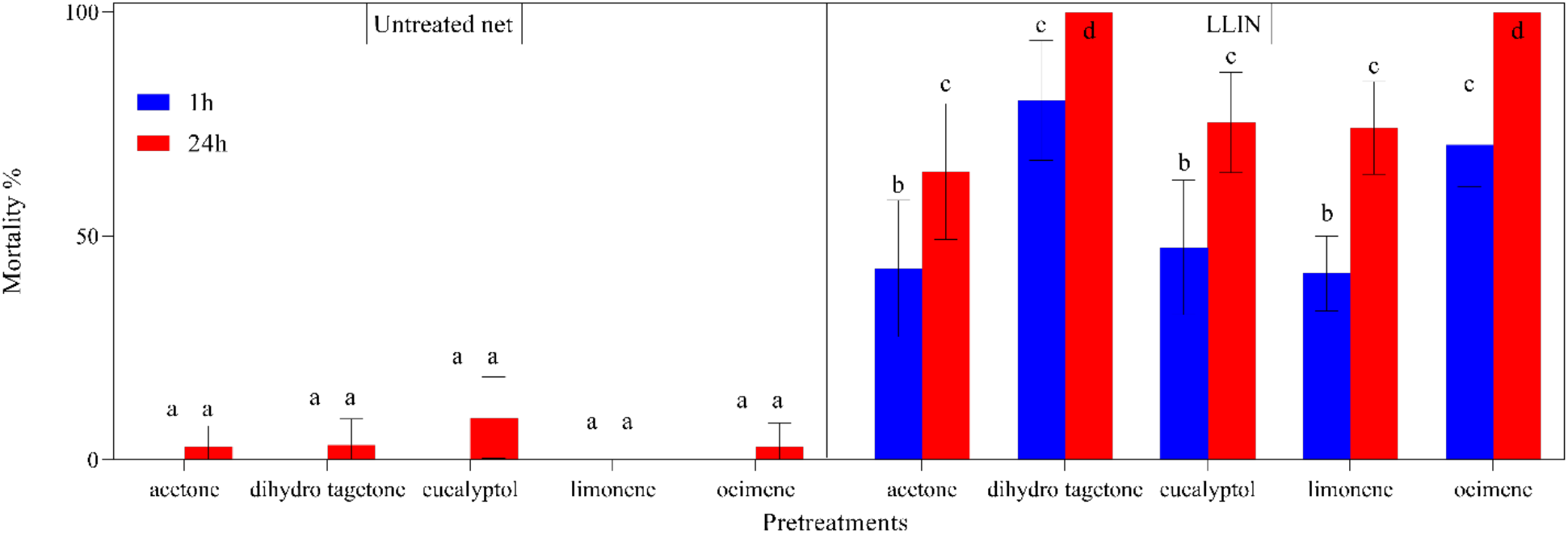
Mortality percentages of 3-5 d old adult female Puerto Rico (*Aedes aegypti*) mosquitoes exposed to oil isolate or acetone (control) vapors for 4 h-pretreatments and transferred briefly to either an untreated net (control, left), or a LLIN (right). Mortality was assessed after 1 h (blue) and 24 h (red) post-contact with the nets. Data presented as mean mortality percentages ± standard error for 60-120 mosquitoes (10 mosquitoes per replicate, number of replicates varies per treatment due to repetition of certain conditions). Statistical differences were obtained with a three-way ANOVA and Tukey’s multiple comparison test, statistical differences are indicated by different letters.

## Discussion

We previously showed that essential oil vapors could affect pyrethroid efficacy after insecticide topical application. Here, we show that, not only do select essential oil vapors affect efficacy of a LLIN after a brief contact, but also that some isolated compounds found in these oils can synergize the effects of an LLIN as well. Amyris oil decreases LLIN efficacy, which was expected based on previous findings (O’Neal et al. 2019), while tagetes and cajeput vapors both increase mortality after contact with the net. The experimental design attempted to mimic a short landing time on the net similar to that in the field. The deltamethrin control (with acetone vapors) caused under 50% mortality, showing some efficacy of the net even after a short contact. The mortality effects of the vapor pre-treatment followed by a brief exposure to the net, were found to be fast acting because mortality was observed within 1 h, and mosquitoes did not recover after 24 h. The main components of the essential oils tested were dihydro tagetone, eucalyptol, D-limonene, and ocimene, but only dihydro tagetone and ocimene had a significant effect on mortality by synergizing the effects of the net within 1 h, an observation that was confirmed after 24 h. It is interesting to note that both dihydro tagetone and ocimene were chosen after GC-MS analysis as components of tagetes oil, and that the other two compounds eucalyptol and D-limonene do not increase mortality on their own, which is surprising due to their significant presence in cajeput oil. This may indicate that these components are not responsible on their own for the effects observed with cajeput. However, the combination of multiple compounds may be responsible for the observed effects, which remains to be tested.

The synergistic potential of the oils and isolates tested with deltamethrin are particularly critical due to the use of a pyrethroid-resistant strain of *Ae. aegypti*, PR, exhibiting multiple resistance mechanisms, including increased P450 activity and expression as well as mutations in the voltage-gated sodium channel (Estep et al. 2017, Rault et al. 2019). We previously showed that the synergistic vapors of essential oils, particularly of tagetes, could reduce P450 activity (O’Neal et al. 2019). This decreased detoxification would circumvent an important resistance mechanism in PR and make deltamethrin efficacious to pyrethroid-resistant mosquitoes. This could have important implications if the method were to be developed for the control of multiple strains and species, which may affect the outcome depending on their resistance mechanism(s). It was shown, however, that many populations of different mosquito species exhibit increased P450 activity as a resistance mechanism (Komagata et al. 2010, David et al. 2013, Dusfour et al. 2015). It would also be interesting to compare the results obtained with a different species exhibiting pyrethroid-resistance, such as a species of the *Anopheles* genus, since malaria mosquitoes are typically targeted using LLINs (Center for Disease Control and Prevention 2019).

These results also confirm that the choice of essential oil is critical. As we have previously shown, only a few synergize deltamethrin as vapor pre-treatments (O’Neal et al. 2019), but some of their separate components such as dihydro tagetone and ocimene may be as or more efficient at facilitating high mortality rates within 1 h. Very little research has been done on dihydro tagetone, but ocimene is a compound commonly found in plants that has been shown to indirectly increase the ability of plants to protect themselves against herbivorous insect infestations (Cascone et al. 2015). However, this compound is usually not delivered as vapor. Using natural components, such as ocimene, in combination with deltamethrin-treated nets can offer more options for synergists in addition to piperonyl butoxide (PBO), which is currently used in some commercialized nets like Permanet 3.0^®^ (Vestergaard 2022). This study highlights two compounds that could be used in a vaporous emanation device to offer additional protection and restored efficacy of pyrethroid-treated nets in the fight against community transmission of mosquito-borne pathogens that lead to human diseases.

## Acknowledgments

We thank Xi Xian Ng for his technical assistance with maintaining the mosquito colonies.

## References cited

Abbott, W. S. 1925. A Method of Computing the Effectiveness of an Insecticide. J. Econ. Entomol. 18: 265–267.

Balabanidou, V., A. Kampouraki, M. MacLean, G. J. Blomquist, C. Tittiger, M. P. Juárez, S. J. Mijailovsky, G. Chalepakis, A. Anthousi, A. Lynd, S. Antoine, J. Hemingway, H. Ranson, G. J. Lycett, and J. Vontas. 2016. Cytochrome P450 associated with insecticide resistance catalyzes cuticular hydrocarbon production in Anopheles gambiae. Proc. Natl. Acad. Sci. 113: 9268–9273.

Brengues, C., N. J. Hawkes, F. Chandre, L. McCarroll, S. Duchon, P. Guillet, S. Manguin, J. C. Morgan, and J. Hemingway. 2003. Pyrethroid and DDT cross-resistance in Aedes aegypti is correlated with novel mutations in the voltage-gated sodium channel gene. Med. Vet. Entomol. 17: 87–94.

Caraballo, H., and K. King. 2014. Emergency department management of mosquito-borne illness: malaria, dengue, and West Nile virus. Emerg. Med. Pract. 16: 1–23; quiz 23–24.

Cascone, P., L. Iodice, M. E. Maffei, S. Bossi, G. Arimura, and E. Guerrieri. 2015. Tobacco overexpressing β-ocimene induces direct and indirect responses against aphids in receiver tomato plants. J. Plant Physiol. 173: 28–32.

Center for Disease Control and Prevention. 2019. CDC - Malaria - Malaria Worldwide - How Can Malaria Cases and Deaths Be Reduced? - Insecticide-Treated Bed Nets. (https://www.cdc.gov/malaria/malaria_worldwide/reduction/itn.html).

Che-Mendoza, A., A. Medina-Barreiro, E. Koyoc-Cardeña, V. Uc-Puc, Y. Contreras-Perera, J. Herrera-Bojórquez, F. Dzul-Manzanilla, F. Correa-Morales, H. Ranson, A. Lenhart, P. J. McCall, A. Kroeger, G. Vazquez-Prokopec, and P. Manrique-Saide. 2018. House screening with insecticide-treated netting provides sustained reductions in domestic populations of Aedes aegypti in Merida, Mexico. PLoS Negl. Trop. Dis. 12: e0006283.

David, J.-P., H. M. Ismail, A. Chandor-Proust, and M. J. I. Paine. 2013. Role of cytochrome P450s in insecticide resistance: impact on the control of mosquito-borne diseases and use of insecticides on Earth. Philos. Trans. R. Soc. B Biol. Sci. 368.

Dery, D. B., G. K. Ketoh, J. Chabi, G. Apetogbo, I. A. Glitho, T. Baldet, and J.-M. Hougard. 2012. Efficacy of a Mosaic Long-Lasting Insecticide Net, PermaNet 3.0, against Wild Populations of Culex quinquefasciatus in Experimental Huts in Togo. ISRN Infect. Dis. 2013: e209654.

Du, Y., Y. Nomura, B. S. Zhorov, and K. Dong. 2016. Sodium Channel Mutations and Pyrethroid Resistance in Aedes aegypti. Insects. 7.

Dusfour, I., P. Zorrilla, A. Guidez, J. Issaly, R. Girod, L. Guillaumot, C. Robello, and C. Strode. 2015. Deltamethrin Resistance Mechanisms in Aedes aegypti Populations from Three French Overseas Territories Worldwide. PLoS Negl. Trop. Dis. 9: e0004226.

Estep, A. S., N. D. Sanscrainte, C. M. Waits, J. E. Louton, and J. J. Becnel. 2017. Resistance Status and Resistance Mechanisms in a Strain of Aedes aegypti (Diptera: Culicidae) From Puerto Rico. J. Med. Entomol. 54: 1643–1648.

Francis, S., K. Saavedra-Rodriguez, R. Perera, M. Paine, W. C. Black, and R. Delgoda. 2017. Insecticide resistance to permethrin and malathion and associated mechanisms in Aedes aegypti mosquitoes from St. Andrew Jamaica. PLOS ONE. 12: e0179673.

Franklinos, L. H. V., K. E. Jones, D. W. Redding, and I. Abubakar. 2019. The effect of global change on mosquito-borne disease. Lancet Infect. Dis. 19: e302–e312.

Gammon, D. W., M. A. Brown, and J. E. Casida. 1981. Two classes of pyrethroid action in the cockroach. Pestic. Biochem. Physiol. 15: 181–191.

Ghosh, D., A. Alim, M. M. Huda, C. M. Halleux, M. Almahmud, P. L. Olliaro, G. Matlashewski, A. Kroeger, and D. Mondal. 2021. Comparison of Novel Sandfly Control Interventions: A Pilot Study in Bangladesh. Am. J. Trop. Med. Hyg. 105: 1786–1794.

GÖKÇE, A., G. Bingham, and M. Whalon. 2018. Impact of long-lasting insecticide-incorporated screens on Colorado potato beetle and plum curculio. Turk. J. Agric. For. 42: 38–44.

Hemingway, J., B. J. Beaty, M. Rowland, T. W. Scott, and B. L. Sharp. 2006. The Innovative Vector Control Consortium: improved control of mosquito-borne diseases. Trends Parasitol., Practical parasitology. 22: 308–312.

Hemingway, J., R. G. Boddington, J. Harris, and S. J. Dunbar. 1989. Mechanisms of insecticide resistance in Aedes aegypti (L.) (Diptera: Culicidae) from Puerto Rico. Bull. Entomol. Res. 79: 123–130.

Hemingway, J., and H. Ranson. 2000. Insecticide resistance in insect vectors of human disease. Annu. Rev. Entomol. 45: 371–391.

Herrera-Bojórquez, J., E. Trujillo-Peña, J. Vadillo-Sánchez, M. Riestra-Morales, A. Che-Mendoza, H. Delfín-González, N. Pavía-Ruz, J. Arredondo-Jimenez, E. Santamaría, A. E. Flores-Suárez, G. Vazquez-Prokopec, and P. Manrique-Saide. 2020. Efficacy of Long-lasting Insecticidal Nets With Declining Physical and Chemical Integrity on Aedes aegypti (Diptera: Culicidae). J. Med. Entomol. 57: 503–510.

Hougard, J.-M., S. Duchon, F. Darriet, M. Zaim, C. Rogier, and P. Guillet. 2003. Comparative performances, under laboratory conditions, of seven pyrethroid insecticides used for impregnation of mosquito nets. Bull. World Health Organ. 81: 324–333.

Johnson, E. 2021. Examination of Cajeput Oil (Melaleuca cajuputi) Phytochemicals as Tools to Manage the Yellow Fever Mosquito (Aedes aegypti L.). Diss. Stud. Res. Entomol.

Komagata, O., S. Kasai, and T. Tomita. 2010. Overexpression of cytochrome P450 genes in pyrethroid-resistant Culex quinquefasciatus. Insect Biochem. Mol. Biol. 40: 146–152.

Kyle, J. L., and E. Harris. 2008. Global Spread and Persistence of Dengue. Annu. Rev. Microbiol. 62: 71–92.

Lumjuan, N., L. McCarroll, L. Prapanthadara, J. Hemingway, and H. Ranson. 2005. Elevated activity of an Epsilon class glutathione transferase confers DDT resistance in the dengue vector, Aedes aegypti. Insect Biochem. Mol. Biol. 35: 861–871.

Manjarres-Suarez, A., and J. Olivero-Verbel. 2013. Chemical control of Aedes aegypti: a historical perspective. Rev. Costarric. Salud Pública. 22: 68–75.

Muktar, Y., N. Tamerat, and A. Shewafera. 2016. Aedes aegypti as a Vector of Flavivirus. J. Trop. Dis. Public Health. 4.

O’Neal, S. T., E. J. Johnson, L. C. Rault, and T. D. Anderson. 2019. Vapor delivery of plant essential oils alters pyrethroid efficacy and detoxification enzyme activity in mosquitoes. Pestic. Biochem. Physiol. 157: 88–98.

Powers, A. M., and C. H. Logue. 2007. Changing patterns of chikungunya virus: re-emergence of a zoonotic arbovirus. J. Gen. Virol. 88: 2363–2377.

Ranson, H., J. Burhani, N. Lumjuan, and W. C. I. Black. 2010. Insecticide resistance in dengue vectors. Trop. Online. 1.

Ranson, H., R. N’Guessan, J. Lines, N. Moiroux, Z. Nkuni, and V. Corbel. 2011. Pyrethroid resistance in African anopheline mosquitoes: what are the implications for malaria control? Trends Parasitol. 27: 91–98.

Rault, L. C., S. T. O’Neal, E. J. Johnson, and T. D. Anderson. 2019. Association of age, sex, and pyrethroid resistance status on survival and cytochrome P450 gene expression in Aedes aegypti (L.). Pestic. Biochem. Physiol. 156: 96–104.

Smith, L. B., S. Kasai, and J. G. Scott. 2016. Pyrethroid resistance in Aedes aegypti and Aedes albopictus: Important mosquito vectors of human diseases. Pestic. Biochem. Physiol. 133: 1–12.

Soderlund, D. M. 2012. Molecular Mechanisms of Pyrethroid Insecticide Neurotoxicity: Recent Advances. Arch. Toxicol. 86: 165–181.

Strode, C., C. S. Wondji, J.-P. David, N. J. Hawkes, N. Lumjuan, D. R. Nelson, D. R. Drane, S. H. P. P. Karunaratne, J. Hemingway, W. C. Black, and H. Ranson. 2008. Genomic analysis of detoxification genes in the mosquito Aedes aegypti. Insect Biochem. Mol. Biol. 38: 113–123.

Thompson, M., J. C. Steichen, and R. H. Ffrench-Constant. 1993. Conservation of cyclodiene insecticide resistance-associated mutations in insects. Insect Mol. Biol. 2: 149–154.

Vestergaard. 2022. PermaNet® 3.0. Vestergaard.

Vontas, J., E. Kioulos, N. Pavlidi, E. Morou, A. della Torre, and H. Ranson. 2012. Insecticide resistance in the major dengue vectors Aedes albopictus and Aedes aegypti. Pestic. Biochem. Physiol., Special Issue: Molecular Approaches to Pest Control, Toxicology and Resistance. 104: 126–131.

Wood, O. R., S. Hanrahan, M. Coetzee, L. L. Koekemoer, and B. D. Brooke. 2010. Cuticle thickening associated with pyrethroid resistance in the major malaria vector Anopheles funestus. Parasit. Vectors. 3: 67.

World Health Organization. 2022. Vector-borne diseases. (https://www.who.int/news-room/fact-sheets/detail/vector-borne-diseases).

Zaim, M., A. Aitio, and N. Nakashima. 2000. Safety of pyrethroid-treated mosquito nets. Med. Vet. Entomol. 14: 1–5.

Zar, J. H. 2010. Biostatistical Analysis, Illustrated, 5th. ed. Prentice Hall.

